# An alarm system for biomedical construct design: a lesson from the unintended protein product of eGFP

**DOI:** 10.64898/2026.06.02.729456

**Authors:** Ziyi Wang, Hongyi Ma, Yafei Mao, Kaiyue Ma

**Affiliations:** Bio-X Institutes, Key Laboratory for the Genetics of Developmental and Neuropsychiatric Disorders, Ministry of Education, International Peace Maternity and Child Health Hospital, Shanghai Key Laboratory of Embryo Original Diseases, School of Medicine, Shanghai Jiao Tong University, Shanghai, China; Zhiyuan College, Shanghai Jiao Tong University, Shanghai, China; Center for Genomic Research, International Institutes of Medicine, Fourth Affiliated Hospital, Zhejiang University, Yiwu, China; Center for Comparative Biomedicine, Ministry of Education Key Laboratory of Systems Biomedicine, State Key Laboratory of Medical Genomics, Institute of Systems Biomedicine, Shanghai Jiao Tong University, Shanghai, China

**Keywords:** plasmid, open reading frame, hidden coding potential, eGFP, gene expression

## Abstract

Plasmids are widely used for gene expression, yet their coding potential beyond the intended coding sequence (CDS) is often poorly characterized. Here, we explored putative “hidden open reading frames” (hidden ORFs) embedded within non-canonical reading frames of plasmid sequences through a computational workflow for their identification. Using enhanced green fluorescent protein (eGFP) as a target gene, we observed unexpectedly uninterrupted ORFs in both the +2 coding frame and the reverse frame. Immunoblotting detected stable expression of the +2 frame-derived protein, but not the reverse-frame ORF. Motivated by these observations, we developed a computational pipeline and analyzed 6,308 eGFP-containing plasmids, identifying putative hidden ORFs in approximately 6% of constructs. Approximately 94% of hidden ORFs occurred in the +2 frame, with the remainder occurring in the reverse frame. The same analytical pipeline, if utilized for plasmids beyond eGFP plasmids, can contribute to avoiding unintended outcomes, in applications such as gene replacement therapy.

---

In light of recent dark proteome studies highlighting widespread translation from non-canonical open reading frames (ncORFs) [1], we realized that similar hidden coding potential may also exist in artificial plasmids and potentially interfere with experiments. An example is the commonly used eGFP. Examination of the eGFP DNA sequence across all six possible translation frames revealed an unexpected feature. For simplicity, we designated the intended coding frame as the canonical frame (CF), with alternative forward frames denoted CF+1 and CF+2 and reverse-strand frames designated RF, RF+1 and RF+2 (Fig. 1a). While the CF encodes GFP, both the CF+2 and the RF contained long uninterrupted ORFs lacking premature stop codons. We refer to such alternative-frame ORFs outside the intended coding frame as hidden ORFs. By contrast, the ancestral GFP from *Aequorea victoria* (avGFP) sequence did not exhibit similarly extended ORFs in alternative frames (Fig. 1b), suggesting that these hidden coding features may have emerged during the engineering and optimization process that generated eGFP.

**Fig. 1.**
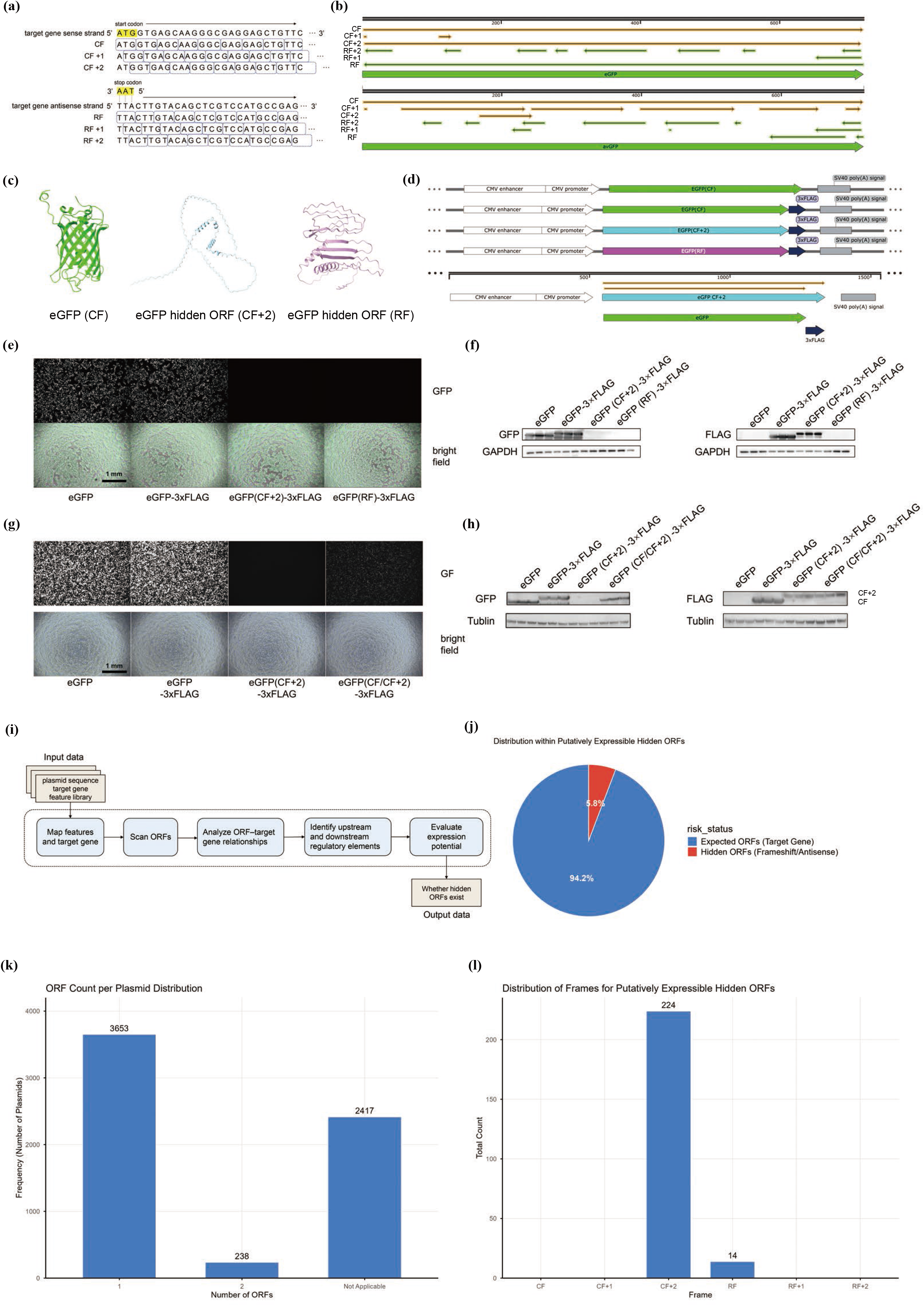
(a) Schematic representation of the six possible translation frames within the eGFP coding sequence. The intended coding frame is defined as the canonical frame (CF), with alternative forward frames designated CF+1 and CF+2 and reverse-strand frames designated RF, RF+1, and RF+2. (b) Comparison of ORF distributions across all six translation frames between eGFP and *Aequorea victoria* GFP (*av*GFP). (c) AlphaFold3-predicted structures of translation products derived from the CF, CF+2, and RF frames of eGFP. (d) Schematic plasmid map containing the eGFP, eGFP-3×FLAG, eGFP(CF+2)-3×FLAG, eGFP(RF)-3×FLAG or eGFP(CF/CF+2)-3×FLAG, arranged from top to bottom. In the eGFP(CF/CF+2)-3×FLAG construct, the 3×FLAG tag is fused in frame with the CF+2 ORF rather than the CF ORF. (e) Fluorescence microscopy analysis of HEK293T cells transfected with eGFP, eGFP-3×FLAG, eGFP(CF+2)-3×FLAG or eGFP(RF)-3×FLAG constructs. Scale bars, 1 mm. (f) Immunoblot analysis of proteins expressed from eGFP, eGFP-3×FLAG, eGFP(CF+2)-3×FLAG or eGFP(RF)-3×FLAG constructs using anti-GFP and anti-FLAG antibodies. GAPDH is served as a loading control. (g) Fluorescence microscopy analysis of HEK293T cells transfected with eGFP, eGFP-3×FLAG, eGFP(CF+2)-3×FLAG or eGFP(CF/CF+2)-3×FLAG constructs. Scale bars, 1 mm. (h) Immunoblot analysis of proteins expressed from eGFP, eGFP-3×FLAG, eGFP(CF+2)-3×FLAG or eGFP(CF/CF+2)-3×FLAG constructs using anti-GFP and anti-FLAG antibodies. β-tublin is served as a loading control. (i) Overview of the computational pipeline developed for hidden ORF identification. (j) Frequency of putatively expressible hidden ORFs identified from 6,308 eGFP-containing plasmids. (k) Distribution of the number of ORFs detected per plasmid. (l) Distribution of putatively expressible hidden ORFs across alternative translation frames.

To determine whether these hidden ORFs might encode distinct proteins, we first analyzed the predicted translation products using *AlphaFold3*. Structural predictions revealed markedly different architectures among the CF, CF+2, and RF translation products (Fig. 1c). We next constructed eGFP, eGFP-3×FLAG, eGFP(CF+2)-3×FLAG, and eGFP(RF)-3×FLAG expression plasmids and transfected them into HEK293T cells for experimental validation (Fig. 1d). Fluorescence microscopy showed that both CF+2 and RF constructs lacked detectable green fluorescence (Fig. 1e). Immunoblotting further detected stable expression of the CF+2-derived protein at the expected molecular weight, but not the RF product (Fig. 1f), demonstrating that at least a subset of hidden ORFs can generate unintended stable protein products.

To determine whether the CF+2 ORF can be translated in the context of the native eGFP coding sequence, we generated eGFP(CF/CF+2)-3×FLAG plasmid, a construct containing both the CF and the CF+2 ORF, with an in-frame CF+2 ATG introduced upstream of the CF+2 ORF while preserving the canonical eGFP ORF (Fig. 1d). A 3×FLAG tag was inserted in frame with the CF+2 ORF to specifically detect its translation. HEK293T cells were transiently transfected with eGFP (CF), eGFP (CF)-3×FLAG, CF+2-3×FLAG, or eGFP(CF/CF+2)-3×FLAG. As expected, GFP fluorescence was detected only in constructs containing the canonical eGFP ORF. Compared with the canonical construct, eGFP(CF/CF+2)-3×FLAG displayed reduced GFP fluorescence (Fig. 1g), suggesting that translation of the CF+2 ORF may partially affect canonical eGFP expression. Western blot analysis using an anti-FLAG antibody detected the expected CF+2-derived protein in cells expressing eGFP(CF/CF+2)-3×FLAG, demonstrating that the hidden CF+2 ORF can be stably translated in the context of the native eGFP coding sequence. Cells transfected with eGFP(CF/CF+2)-3×FLAG exhibited relatively weaker fluorescence signal; however, this reduction was not reflected in our immunoblotting experiment (Fig. 1h), likely owing to the limited quantitative resolution under the specific immunoblotting experimental conditions.

Motivated by this observation, we asked whether hidden ORFs are common within plasmid constructs. We therefore developed a computational pipeline to systematically identify potentially expressible hidden ORFs from plasmid sequences (Fig. 1i). Briefly, plasmid sequences were scanned across all six translation frames on both strands to identify ATG-initiated ORFs exceeding 100 amino acids. Candidate ORFs were annotated according to their positional relationship with the target gene and evaluated for the presence and proximity of upstream promoter and downstream transcriptional termination elements. ORFs located in non-canonical frames and associated with these transcriptional features were classified as putatively expressible hidden ORFs.

Applying this framework to 6,308 eGFP-containing plasmids from Addgene revealed putatively expressible hidden ORFs in approximately 6% of constructs (Fig. 1j). A subset of plasmids was classified as Not Applicable because no mammalian transcriptional regulatory elements could be identified based on our curated feature library or because they lacked mammalian expression cassettes. Among the remaining plasmids, approximately 94% contained only the canonical eGFP ORF, whereas about 6% harbored one additional hidden ORF. (Fig. 1k). Approximately 94% of putatively expressible hidden ORFs occurred in the CF+2 frame, whereas the remainder originated from the reverse frame (Fig. 1l), consistent with our initial observation that eGFP harbors long uninterrupted ORFs specifically in these alternative frames. No hidden ORFs were identified in other translation frames, due to the frequent occurrence of premature stop codons that interrupt long ORF formation. These findings suggest that hidden coding potential may be more prevalent than previously appreciated in commonly used plasmid constructs.

The existence of hidden ORFs may have implications extending beyond routine plasmid experiments. First, translation of hidden ORFs could potentially influence expression of the intended protein product, as internal alternative ORFs have been reported to modulate translation of their corresponding main ORFs in bacterial systems [2]. Second, hidden ORFs may give rise to unintended protein products with biological activities distinct from those of the encoded transgene. For example, proteogenomic and immunological studies have shown that peptides derived from non-canonical reading frames can be translated and enter antigen-presentation pathways, including MHC-I-mediated immune recognition [3,4]. In therapeutic contexts such as gene replacement therapy, hidden ORFs may therefore represent a previously overlooked source of unintended protein expression, which could potentially generate unanticipated biological effects, including immune responses and potential toxicity. Although the functional consequences of most hidden ORFs remain unknown, our work establishes a framework for systematically identifying hidden coding potential within plasmid sequences and highlights its potential relevance for the design and safety evaluation of the constructs for gene replacement therapy.

## Acknowledgements

We thank the Addgene team (Dr. Jason Niehaus, Dr. Hannah Dotson, and Dr. Chonnettia Jones) for providing plasmid sequence datasets used in this study. This work is supported by SJTU Zhiyuan Honorary Scholarship Program to Z.W. This work is in part supported by the Shanghai Rising-Star Program (24YF2721800), the Shanghai Magnolia Talent Plan Pujiang Project (24PJA049), and the Shanghai Post-doctoral Excellence Program (2024338) to K.M. This work is in part supported by the Scientific Research Innovation Capability Support Project for Young Faculty (SRICSPYF-ZY2025101), the National Key Research and Development Program of China (2025YFC3410300), the National Natural Science Foundation of China (32370658), the Natural Science Foundation of Chongqing, China (CSTB2024NSCQ-JQX0004), the New Cornerstone Science Foundation through the XPLORER PRIZE, Shanghai Jiao Tong University (SJTU) 2030 Initiative (WH510363003/016), Yongxin Youth Award Fund and Zhongying Young Scholars Program to Y.M.

## CRediT authorship contribution statement

**Ziyi Wang:** Methodology, Software, Formal analysis, Investigation, Writing - Original Draft, Visualization, Data Curation, Funding acquisition. **Hongyi Ma:** Investigation, Writing - Review & Editing. **Yafei Mao:** Resources, Writing - Review & Editing, Funding acquisition. **Kaiyue Ma:** Conceptualization, Methodology, Investigation, Resources, Writing - Review & Editing, Supervision, Project administration, Funding acquisition.

## Supplementary Materials

Supplementary Materials are available online.

## Data availability

The computational pipeline developed in this study is publicly available at: https://github.com/ZiyiWang-sjtu/hidden-ORFs-search. Plasmid sequence data were provided by the Addgene team. Processed datasets generated during this study are available from the corresponding author upon reasonable request.

