## Supplementary material for "An alarm system for biomedical construct design: a lesson from the unintended protein product of eGFP": Supplemental_Material.docx

**Supplementary methods**

**Plasmid construction**

The plasmids were constructed by inserting CMV-eGFP, CMV-eGFP-3×FLAG, CMV-eGFP(CF+2)-3×FLAG, and CMV-eGFP(RF)-3×FLAG into the pMPRA1 plasmid (MiaoLingBio, P31645) between the pause site and the SV40 poly(A) signal, respectively. The cloning of the CMV-eGFP plasmid was previously documented [1]. The CMV-eGFP(CF/CF+2)-3×FLAG plasmid was generated by Gibson assembly. The backbone was obtained by NcoI (NEB, #R0193S) digestion of the CMV-eGFP(CF+2)-3×FLAG plasmid, while the insert was PCR-amplified from the same plasmid using primers. PCR was performed using Q5 High-Fidelity DNA Polymerase (NEB, #M0491L). NEBuilder HiFi DNA Assembly Master Mix (NEB, #E2621S) was used for the assembly. DH5α chemically competent cells (Vazyme, #C502) were used for the transformation. The plasmids were sequenced and confirmed with Sanger sequencing (BioSune) and whole plasmid sequencing. The primers used and the plasmid sequence are listed in the supplementary table.

**Cell culture and transfections**

HEK293T cells were maintained in Dulbecco’s Modified Eagle’s Medium (DMEM; Servicebio, #G4515) supplemented with 10% fetal bovine serum (FBS; Servicebio, #G8003), 1× Penicillin-Streptomycin (Servicebio, #G4003), at 37 °C with 5% CO2. Transfection was performed using Lipomaster 3000 Transfection Reagent (Vazyme, #TL301) and Opti-MEM I Reduced Serum Medium (Gibco, #31985062).

**Immunoblotting**

Cells were harvested and lysed in RIPA buffer (Sigma-Aldrich, #R0278-500mL) supplemented with protease inhibitor. Protein concentrations were determined using a BCA protein assay kit (Servicebio, #G2026-1000T). Equal amounts of protein were mixed with SDS loading buffer (YEASEN, #20315ES05), denatured at 95°C for 5 min, separated by SDS–PAGE (Genefist, #GF1820-08), and transferred onto PVDF membranes (Bio-rad, #1620264;).

Membranes were blocked in Fast Blocking Buffer (Servicebio, #G2052-500ML) and incubated with primary antibodies against GFP (Invitrogen, #A6455), FLAG (Merck, #F1804-50UG), or overnight at 4°C. After washing, membranes were incubated with the corresponding HRP-conjugated anti-mouse (Servicebio, #GB23301) or anti-rabbit (Servicebio, #GB23303) secondary antibodies and visualized using enhanced chemiluminescence (Servicebio, #G2014-500ML). For experiments using GAPDH as the loading control, membranes were subsequently stripped with antibody stripping buffer (Servicebio, #G2078-100ML), incubated overnight at 4°C with an anti-GAPDH antibody (Proteintech, #81640-5-RR), followed by incubation with the corresponding HRP-conjugated secondary antibody and chemiluminescent detection. For experiments using β-tubulin as the loading control, membranes were incubated overnight at 4°C with an anti-β-tubulin antibody (huabio, #ET1602-4), followed by incubation with the corresponding HRP-conjugated secondary antibody and chemiluminescent detection.

**Computational pipeline of hidden ORFs search**

The pipeline takes plasmid sequences, target gene sequences, and annotated regulatory features as input and generates ORF annotations and associated metrics, including reading-frame classifications, positional relationships to the target gene, and putative expression status.

Plasmid sequences were obtained from the Addgene team, which contains sequence records for 156,799 plasmids. Plasmids containing eGFP sequences were extracted from the the addgene-plasmids-sequences.json dataset by sequence matching against the eGFP coding sequence. Regulatory feature annotations were curated from SnapGene records using SnapGene software (http://www.snapgene.com/), including promoters, polyadenylation (polyA) signals, and long terminal repeat (LTR) elements, and compiled into a feature library. Only mammalian regulatory elements were retained for subsequent analyses.

Plasmids were treated as circular molecules by concatenating each sequence to itself, enabling the identification of ORFs or regulatory elements spanning the plasmid origin. The first stage of the pipeline identified hidden ORFs, defined as all ATG-initiated ORFs present at the DNA sequence level regardless of transcriptional regulatory elements. Six-frame translation scanning was then performed across both strands to identify ATG-initiated ORFs exceeding a minimum length threshold of 100 amino acids.

For each identified ORF, positional relationships relative to the target gene were classified as outside, inside, spanning, or partial overlap. Relative reading-frame relationships were defined with respect to the target gene coding sequence and designated as CF, CF+1, CF+2, RF, RF+1, and RF+2.

In the second stage, mammalian regulatory features were identified by BLASTN searches against plasmid sequences using a 96% sequence identity threshold and a minimum coverage of 90% of the feature length, allowing robust feature detection while tolerating minor sequence variations among plasmid records. Distances between each ORF and the nearest upstream promoter and downstream transcriptional termination element (polyA signal or terminator) were then calculated. ORFs were further evaluated according to predefined structural criteria, including a minimum length of 100 amino acids, overlap with the target gene region, a maximum distance of 10 kb from an upstream promoter element, and a maximum distance of 10 kb from a downstream transcriptional termination element. The 10 kb threshold was chosen to accommodate multi-gene expression cassettes rather than to reflect the expected length of a biological untranslated region (UTR). Hidden ORFs located in non-canonical reading frames that satisfied these criteria were classified as putatively expressible hidden ORFs.

The current pipeline relies on BLAST-based annotation of regulatory elements together with distance thresholds to infer putatively expressible hidden ORFs. Although this strategy provides a practical approximation, it may overestimate the expression potential of some reverse-frame hidden ORFs, where the associated promoter or transcriptional termination element can be relatively distant due to the lack of a conventional expression cassette organization. Therefore, the current distance-based criteria should be regarded as a coarse approximation rather than a definitive predictor of expression. More sophisticated computational approaches, such as machine learning or artificial intelligence models, may enable direct identification of functional expression cassettes from plasmid sequences and prediction of the probability and relative strength of hidden ORF expression.

**References:**
1. Fu, L., Chen, J., Lian, D., Du, S., Wu, D., Yang, C., ... & Mao, Y. (2026). A long-read human pangenome initiative for comprehensive interpretation of nuclear-embedded mitochondrial DNA. *Nature Communications, 17*(1), 4371.
