## Supplementary figures and images for "An alarm system for biomedical construct design: a lesson from the unintended protein product of eGFP"

### graphic abstract_PNG.png

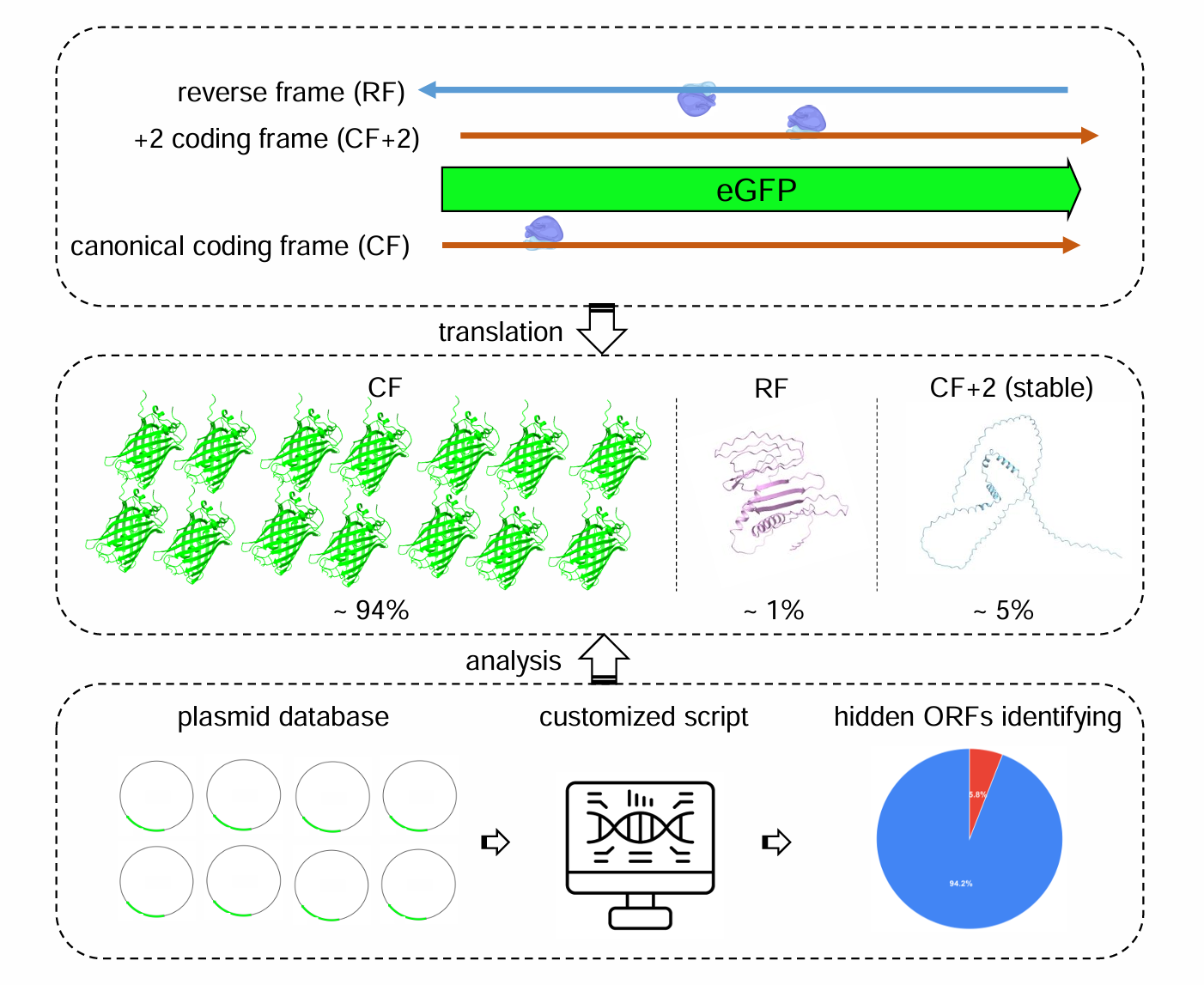
